# When seeps give ANME-SRB the cold shoulder: putative role of denitrification mediated methane oxidation in an Antarctic Cold Seep

**DOI:** 10.64898/2026.08.07.738775

**Authors:** Jacob H. Wynne, Rowan H. McLachlan, Andrew R. Thurber

## Abstract

Antarctica represents a significant, unresolved, and unstable source of methane to the atmosphere. To advance our understanding of the biological filter of methane in Antarctica, here we identify the taxa and functional genes present during methane oxidation in an Antarctic Methane Seep. Methane oxidation was present in all sediments, including in a non seep control site. Using 16S rRNA analysis alongside metagenomics, we found that ANaerobic MEthane oxidizing (ANME) archaea coupled to Sulfate-Reducing Bacteria (SRB), documented as the most important marine methane sink in other locations, were not present. Instead, we observed the presence of denitrification-dependent methane oxidizers, including the anaerobic genus *Candidatus Methylomirabilis*, alongside the nitrate reducing archaea *Candidatus Methanoperedens* through short-read metagenomic classification. In addition, we note the presence of multiple aerobic methanotrophs, with a particularly high abundance of the *Methylobacter*, *Methylomonas*, and *Methyloprofundus* genera. Our results support denitrification-mediated methane oxidation and aerobic methanotrophy as the primary potential methane sinks in the Ross Sea. The widespread methane oxidation, including in ‘control’ sediment, combined with the possibility of anaerobic methane oxidation linked to denitrification rather than sulfate reduction highlights the ubiquity and uniqueness of the Antarctic methane cycle.

## Introduction

Methane is a potent greenhouse gas that is responsible for 20% of atmospheric radiative forcing since 1750 (Jackson et al. 2021; Masson-Delmotte et al. 2021). Concerningly, there has been an increase in methane emissions since 2014 with no known cause (Nisbet et al. 2019). Antarctica represents a vast potential methane reservoir, holding up to 20% of global methane hydrate reservoirs (Ruppel and Kessler 2017; Wadham et al. 2012), and its shallow Gas Hydrate Stability Zone (GHSZ) makes Antarctic hydrate reservoirs one of the most susceptible to climate change (Kretschmer et al. 2015; Giustiniani et al. 2018). Recent findings support this susceptibility, with the emergence of dozens of seafloor seeps in coastal waters of the Ross Sea (Seabrook et al. 2025; Thurber, Seabrook, and Welsh 2020). Therefore, understanding the sinks of marine methane in Antarctica is a critical research question to understand the Earth system in a time of change.

Despite the oceans containing vast reservoirs of methane, the contribution of marine methane to the atmosphere, and thus greenhouse forcing, is low, at <3% annually (Saunois et al. 2025). This is primarily due to the anaerobic oxidation of methane (AOM), a microbial metabolic process that consumes 70-90% of methane produced in marine sediments prior to the methane’s release to the water column (Knittel and Boetius 2009; Orphan et al. 2001). In marine systems, AOM is typically performed by a polyphyletic clade of ANaerobic MEthanotroph (ANME) archaea syntrophically coupled to sulfate-reducing bacteria (SRBs; Knittel and Boetius 2009). ANME have uniquely slow doubling times (2-7 months) which indicates a >60-year lag before AOM communities can respond to climate-driven increases in methane flux (Dale et al. 2008; Knittel and Boetius 2009; Girguis, Cozen, and DeLong 2005; Nauhaus et al. 2007). Intriguingly, recent studies have revealed the absence of anaerobic methane-oxidizing communities in certain methane-rich locations (Lapham et al. 2024; Semler and Dekas 2025) and modeling approaches have suggested that seafloor communities will be unable to adapt sufficiently to mitigate climate driven impacts of methane release (Stranne et al. 2022). These studies highlight emergent and significant knowledge gaps in our understanding of the global methane cycle and the ability of the dominant methane sinks to adapt to change: the impact of biological succession, growth, and species-species interactions of sediment microbiomes on methane release, particularly in Antarctica (Thurber, Seabrook, and Welsh 2020; Semler and Dekas 2025; Klasek et al. 2021; Ruppel and Kessler 2017; Joung et al. 2022).

While initial understanding of AOM focused on using sulfate as the electron acceptor, AOM has expanded from the previously discussed ANME-SRB consortia with additional mechanisms mediated by a number of materials and processes (Guerrero-Cruz et al. 2021). The archaea *Candidatus Methanoperedens nitroreducens*, initially known as ANME-2d, was found to perform AOM without a syntrophic partner, using nitrate as its electron acceptor (Haroon et al. 2013; Raghoebarsing et al. 2006). The bacterium *Candidatus Methylomirabilis oxyfera,* from the NC10 phylum, was found to perform aerobic oxidation of methane anaerobically by reducing nitrite and producing intracellular oxygen (Ettwig et al. 2010). Other electron acceptors have been found to mediate AOM as well in recent years, including humic acids (Bai et al. 2019), manganese, and iron (Beal, House, and Orphan 2009). In cold seeps, although SRB-mediated AOM predominates, studies have found that denitrification- mediated AOM may be an overlooked sink (Jing et al. 2020; Jiang et al. 2023).

The understanding of the metabolic reach of aerobic methanotrophs has increased as well, covering micro-oxic to anoxic conditions (Guerrero-Cruz et al. 2021). Certain gammaproteobacterial methanotrophs such as *Methylobacter*, *Methylococcus*, *Methylomonas,* and others, appear to have adaptations to low oxygen conditions, including high-affinity cytochromes (Guerrero-Cruz et al. 2021; Skennerton et al. 2015) and hemerythrin (Rahalkar and Bahulikar 2018). Gammaproteobacterial methanotrophs are often found at oxic/anoxic sediment interfaces, supporting their ability to leverage micro-oxic conditions as a component of their niche (Guerrero-Cruz et al. 2021). Aerobic methanotrophs, particularly *Methylococcales*, are often present in seep sediments globally and are one of the distinguishing factors of seep sediment against background sediment (Ruff et al. 2015). Aerobic methanotrophs are also highly responsive to changes ecosystem-wide including shifts in methane regimes, dividing more quickly and responding orders of magnitude faster than their anaerobic counterparts (Kessler et al. 2011; Leonte et al. 2017).

Previous research in Antarctica has shown the importance of understanding the microbial community and its succession to constrain the potential impact of methane. Thurber et al. (2020) reported the biogeochemical and microbial features of Cinder Cones Seep (CCS) 1 year and 5 years after seepage began in this location. At this site, ANME-1, which performs AOM with sulfate-reducing bacteria, were not found one year after seepage began, then reached a maximum of 4% of the microbial community 5 years later. However, ANME-1 was not expected based on the high sulfate, low temperature (-1.8°C) conditions present as informed by global biogeographic comparisons (Thurber, Seabrook, and Welsh 2020). *Methylococcales*, an aerobic methanotroph, was the other microbial taxa involved in methane oxidation there (Thurber, Seabrook, and Welsh 2020). Here, we used a series of experiments to link realized methane oxidation at CCS to the microbial taxa involved. Through amplicon and metagenomic approaches, we resolve an unexpected suite of methanotrophs in the region.

## Materials & Methods

### Sites

The Cinder Cones Seep (77° 47.998’ S, 166° 40.241’ E; Figure 1), began seeping methane in 2011, as marked by the emergence of a large microbial mat, with opportunistic sediment cores and imagery collected in 2012 (Thurber, Seabrook, and Welsh 2020). Methane concentrations in the sediment increased to 0.7 mM with co-occurring sulfate and sulfide, without a marked sulfate-methane transition (Thurber et al. 2020). Methane flux and seepage continued through 2023 (Seabrook et al. 2025). The McMurdo Intake Jetty (77° 51.069’ S, 166° 39.855’ E) is considered an off-seep site where methane concentrations were below detection limits in 2022 and 2023.

**Figure 1.**
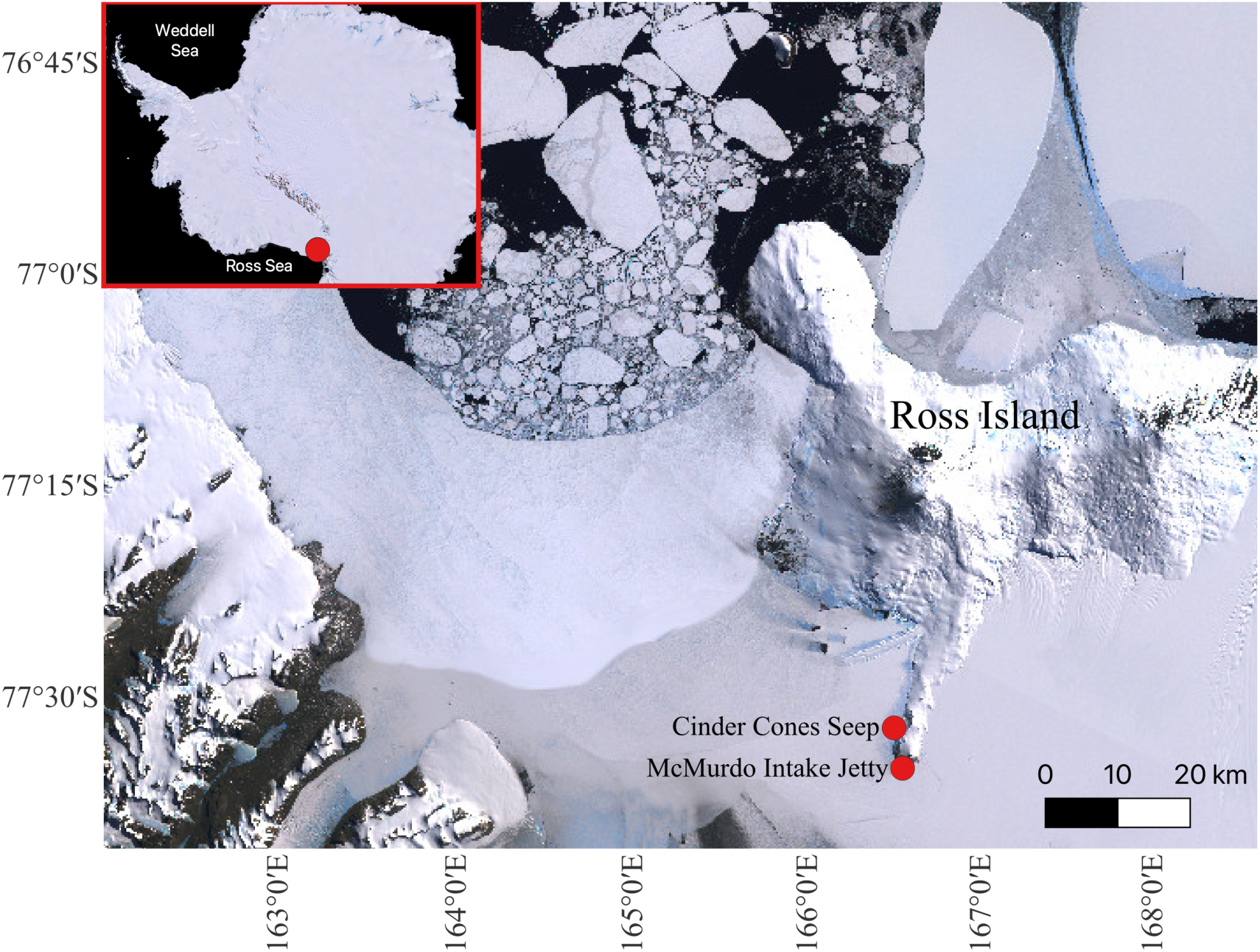
Study Region of the two sites sampled in this study. The map is modified from the Landsat Image Mosaic of Antarctica (LIMA). Unmodified files are freely available through the United States Geological Survey (USGS; https://lima.usgs.gov).

### Sample collection and processing

Our overarching goal was to identify which taxa were present during methane oxidation in both oxic and anoxic conditions, leading to varying incubation times and treatments over the two years of study. Sediment cores (6.4 cm diameter, 10 cm depth) were haphazardly collected from filamentous microbial mats along the CCS in 2022 and 2023 at 10-11m water depth via SCUBA. At the McMurdo intake jetty, cores were taken from non-seep sediments in 2022 at 20m water depth. Each core was immediately placed in a cooler filled with seawater (-1.8°C) and transported to McMurdo station, Antarctica. Samples were kept at *in situ* temperature (-1.8°C) or placed on ice through sample processing and experimentation. The methodology evolved between the years to account for observed potential artifacts based on the mylar bags we used in 2022 (Table S1): these bags allowed oxygen to permeate over time shown through measurements taken at breakdown (Table S2). In 2022, experimental cores were sectioned from 0-3 cm, 3-6 cm, and 6-9 cm and placed in mylar bags. In 2023, experimental cores were sectioned at 0-4 cm and 4-8 cm and placed in glass vials with PTFE-lined caps and kept in the dark.

### Experimental protocol

For experiments 1-4 conducted in 2022, 50 g of sediment was made into a slurry and placed into a mylar bag with 500 mL of 0.22 μm Sterivex-filtered seawater. Incubations were performed with two different treatments: one was injected with 2.5 mL of methane gas isotopically labeled with 30% *δ*^13^*CH*_4_ and 70% *δ*^12^*CH*_4_ while the other incubation served as a no-methane-added control. Sediment was incubated at -1.8°C until methane oxidation was observed (Table S1). Our aim was not to record a quantitative methane oxidation rate, but instead observe when methane oxidation is occurring. During 2 experiments (3 and 4), we ran a FSW control with methane to account for methanotrophs within the water.

For experiments 5 and 6 conducted in 2023, sediment cores were sectioned into 0-4 cm and 4-8 cm depth intervals. In Thurber et al. (2020), this sediment horizon (the transition to 4-5 cm) consistently had taxa with anaerobic metabolisms (including Sulfate reduction, anaerobic methanotrophy, and no aerobic methanotrophs) making anaerobic treatments the most akin to in situ conditions. The 4-8 cm sediment horizon, after initial separation from the 0-4 cm, was processed in an N_2_-flushed glove bag and the water added was sparged with nitrogen to remove oxygen. In contrast to the previous years, experiments were run in 60ml glass vials with PTFE liners; this meant that 10ml of sediment and 50 ml of filtered seawater were added to both oxic (0-4 cm) and anoxic (4-8 cm) treatments and methane volumes were reduced to 0.5 ml for treatments in which it was added. All vials were incubated at -1.8°C for 6 days. At T0 and at the end of each experiment, sediment was preserved in a 1:1 mixture of 2x DNA/RNA shield (Zymo Research, Irvine CA) and stored at -80°C. A filtered seawater control was run in parallel with and without methane added.

We used a unifying naming scheme where samples are labeled by experiment (E), replicate (R) if applicable, depth (e.g., 0_3cm), methane addition (+CH_4_ or -CH_4_), and normoxic (+O2) or anoxic (-O2) water. Full conditions are provided in Table S1.

### Quantifying methane oxidation

Methane oxidation was observed through the movement of ^13^C from methane into the DIC pool as quantified by *Δδ*^13^*C_dic_* via a Picarro G-2131i cavity ring-down spectrometer (CRDS; Picarro, Santa Clara CA). The CRDS was paired with an Automate Prep device connected to a Picarro Liaison for injection and allowed onsite observation of methane oxidation. In brief, 5ml of samples were acidified with 10% phosphoric acid, sparged with nitrogen and the resultant carbon dioxide collected for analysis on the CRDS using the Liaison. For each experiment, *Δδ*^13^*C_dic_* was calculated by taking the difference of *δ*^13^*C_dic_* at the final time point between the methane-added and no-methane-added paired incubations. Values were adjusted across experiments using (Eq 1), and were recorded (Table S3) and visualized (Figure 2).

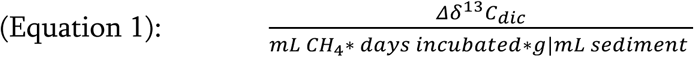

**Figure 2.**
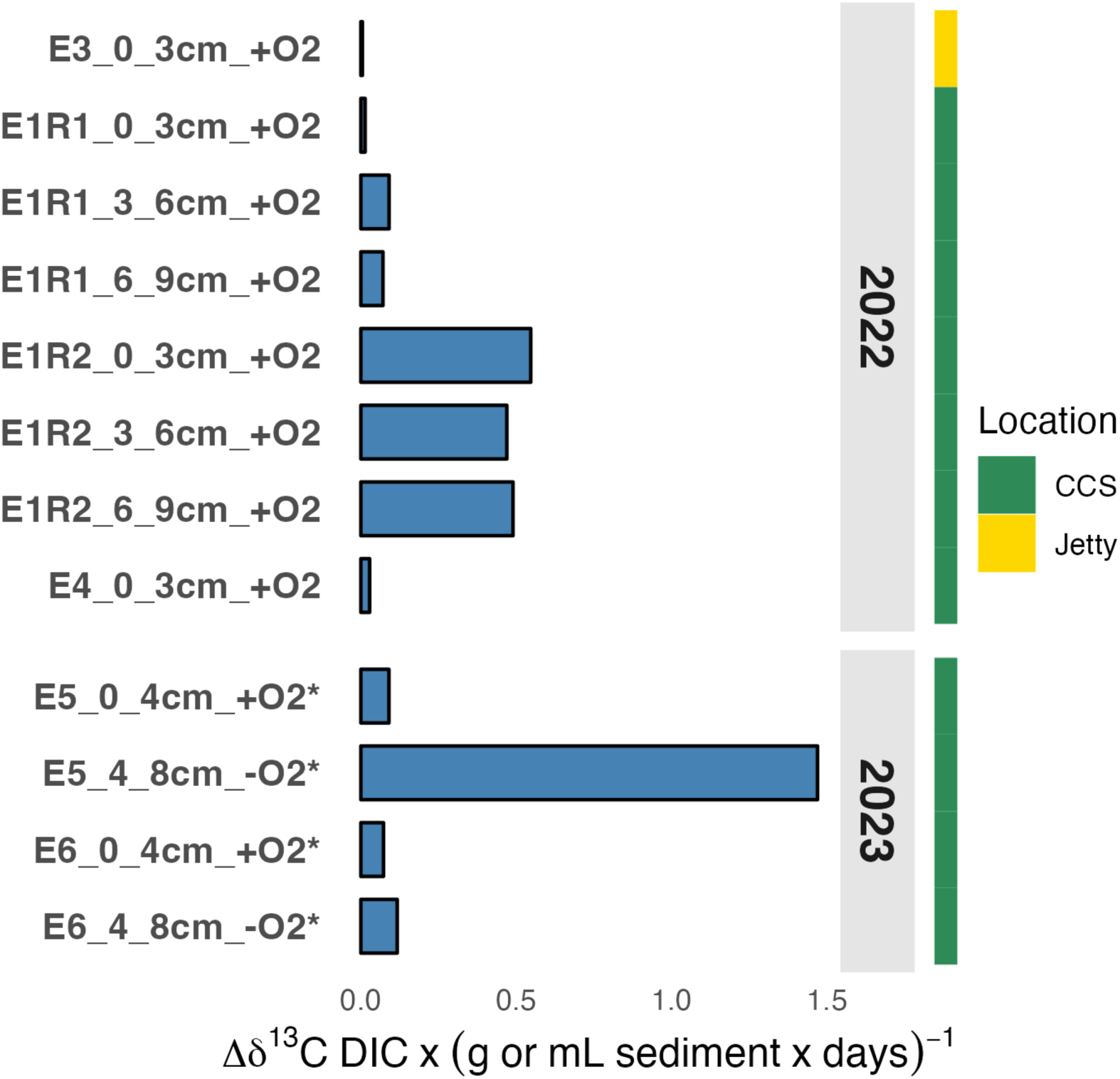
Relative methane oxidation across experimental replicates and treatments. Change in ^13^C_dic_ for each sample incubated with 30% ^13^CH_4_ and 70% ^12^CH_4_, adjusted for amount of methane added, grams (2022) or mL (2023; marked by *) of sediment and number of days.

### DNA extraction and sequencing

DNA was extracted from DNA/RNA shield-preserved samples using the Zymobiomics DNA/RNA miniprep kit (Zymo Research, Irvine CA), and yield was determined using the Qubit 3 fluorometer. For 16S rRNA gene analysis, the Earth Microbiome Project (EMP) protocol was followed (https://github.com/biocore/emp) as employed by Thurber et al. (2020). In brief, PCR amplification was carried out using 515f and 806r primers (Dataset S1), followed by the QIAquick PCR Purification Kit protocol (Qiagen) and sequencing was performed on an Illumina MiSeq (V.2 chemistry and 2 x 250 paired-end sequencing). For metagenomic analysis, library preparation was done at Oregon State University’s Center for Quantitative Life Sciences (CQLS) using the Nextera XT Library Preparation Kit and sequenced on a NextSeq 2000 (2 x 150 bp P3 run).

### Metagenomic taxonomic analysis

Short read taxonomic assignment was carried out using Kaiju with the NCBI refseq_nr database to the genus level (June 6 2023 release; Menzel, Ng, and Krogh 2016). Data were normalized to relative abundance and filtered to exclude taxa and unclassified reads that made up less than 0.001% of the resultant database (including unannotated reads). This cutoff has been suggested for complex microbial communities, yielding high performance while retaining rare taxa within the sample (Edwin et al. 2024). Post filtering, all retained taxonomic assignments had more than 500 reads assigned to them, a threshold for high precision and recall in metagenomic short read classifiers (Buffet-Bataillon et al. 2022).

### Metagenomic community analysis

Metagenomic samples conglomerated at the genus level were analyzed for their differences through depth, experimental approach, and methane exposure. Principal Coordinates Analysis (PCoA), based on Bray-Curtis dissimilarity, was used to visualize differences in community composition (Gower 2015). We tested whether depth, site, methane treatment and oxygen treatment led to differences in the community composition, using a single PERMANOVA with marginal sums of squares using the adonis2 function in vegan (v. 2.6.4) with 999 permutations.

### 16S community and taxonomic analysis

Amplicon Sequence Variants (ASVs) were identified within the 16S rRNA data using Dada2 within the Qiime2 package (Estaki et al. 2020; Bolyen et al. 2019). Taxonomic assignments were carried out through comparison to the Silva v138.1 n99 database which was trimmed to the 515f/806r primer region to improve taxonomic accuracy (Bokulich et al. 2018), using the Qiime Rescript package (Ii et al. 2021) and trained on a naïve Bayes classifier for downstream taxonomy assignments. To reduce the effects of sequencing errors, singleton reads were removed from all samples and the reverse reads (R2) were low quality throughout and omitted from the analytical pipeline. Taxonomic and community analysis were performed on 16S in parallel with metagenomes at the genus level (Figures S1-S4).

### Comparing taxonomic output between 16S and metagenomic methods

Comparison between 16S and metagenomic methods was performed to determine the differences across methods at multiple taxonomic levels. To do this, taxonomically annotated metagenomic reads were filtered as described above to minimize false positives, and singletons were removed from 16S data. The total number of successfully annotated bacterial and archaeal orders, families, and genera were compared between each other to highlight differences across taxonomic boundaries. Potential methanotroph presence/absence was visualized for orders, families and genera containing one or more methanotrophs where presence was called if the taxonomic group was above each method’s respective filtering threshold. To compare methanotrophs consistently across SILVA and NCBI taxonomy, lineages were reconciled and are provided in Tables S4 and S5. For ANME lineages, taxonomy followed (Chadwick et al. 2022). Alongside presence and absence, the total number of reads assigned to each genus were reported within and across samples, alongside the mean, median, and relative abundance metrics (Tables S6, S7).

### Metagenomic functional analysis

Short read functional assignment was carried out to predict the relevant genes related to methane, nitrogen, phosphorus, and sulfur cycling. Trimmed paired-end reads were then merged using PEAR (v 0.9.11; J. Zhang et al. 2014). Merged reads were then annotated by PRODIGAL (--meta option; v 2.6.3; Hyatt et al. 2010) to predict genes, followed by comparison to the databases MCycDB (Qian et al. 2022), PCycDB (Zeng et al. 2022), NCycDB (Tu et al. 2019), and SCycDB (Yu et al. 2021) using DIAMOND (v 2.1.9; Buchfink, Xie, and Huson 2015). We used only the top hit per gene fragment as each gene’s identity. Prior to downstream analysis, DIAMOND alignments were filtered to include hits with an e-value less than 1e^-5^, and a percent identity greater than or equal to 60%.

Annotated genes were grouped into their respective biogeochemical pathways and transformed using the Variance Stabilizing Transformation method within the DESeq2 package (Love, Huber, and Anders 2014). Pathways were visualized using the R package ‘ComplexHeatmap’ (Gu 2022), and row-scaled to z-score so that each pathway is centered on its mean abundance across samples. Distance was measured between samples and functional pathways using Euclidean distance (Danielsson 1980). Clustering between pathways and samples was calculated using the Ward clustering algorithm (Anderberg 1973). To determine the significance of differential abundance of biogeochemical pathways, DESeq2 was used (Love, Huber, and Anders 2014), and data was normalized using its internalized normalization method. Differential abundance testing was performed on methane exposure (yes vs. no), site (Jetty vs. CCS), Year (2022 vs. 2023), and sediment depth. For sediment depth, multiple depths were binned prior to testing between ‘shallow’ (0-3 cm in 2022, 0-4 cm 2023) and ‘deep’ (3-6 cm and 6-9 cm in 2022, 4-8 cm in 2023) depths.

### Analysis code and accessibility

Analysis across the study was conducted via command line using GNU Bash (v. 5.1.8), as well as R (v. 4.2.3) in RStudio (v. 2024.04.1). Portions of the analysis were drafted with assistance from Claude (Anthropic, Opus 4-4.8 and Sonnet 3.5-4.6). All code was reviewed, executed, and validated by the authors, who take full responsibility for its accuracy. All scripts and intermediate files to reproduce results were archived (Wynne, McLachlan & Thurber, 2026).

## Results

### Methane oxidation in sediments

Methane oxidation occurred in all sediment samples that had methane added, however there was high heterogeneity across site, experiments and even replicates (Figure 2). Oxidation ranged from 0.005 *Δδ*^13^*C_dic_*to 1.470 *Δδ*^13^*C_dic_*with a mean of 0.289 *Δδ*^13^*C_dic_*across all samples. The lowest value recorded was by E3_0_3cm_+CH4_+O2, the only experiment conducted on sediment collected from the McMurdo Intake Jetty, an off-seep location, while all other experiments were conducted on sediments collected from CCS. Methane oxidation also occurred regardless of sediment depth. Surprisingly, the only sample different by depth was E5, which had a maximum +1.47 *Δδ*^13^*C_dic_* at 4-8cm which was 16x that of the 0-4cm; this was one of the few samples where anoxic conditions were obtained. Contrastingly, the 2022 E1R2 presented a similar *Δδ*^13^*C_dic_* throughout the sediment horizons (0.06 *Δδ*^13^*C_dic_* across all depth horizons). Upon examination of abiotic controls in 2023, we found no evidence of oxidation present, with negative values from the difference of the methane- added and no-methane-added samples (Table S3). The normoxic filtered-seawater control yielded a -0.11 *Δδ*^13^*C_dic_* while the anoxic filtered-seawater control yielded -0.41 *Δδ*^13^*C_dic_*over the course of the experimental period (Table S3).

### An overview of 16S and metagenomics data

After quality control, the average forward read 16S depth was 126,203 (±55,283) reads with the total number of 16S sequences being 4,290,914 across the whole study (36 samples). Metagenomics yielded an average read depth of 39.3 M (±8.4M) across 34 samples. The total number of reads summed to 2.7 billion across all samples.

### Factors shaping community composition in Antarctic sediment

Depth of sediment and site had a significant impact on the microbial communities within the study while methane and oxygen-exposure did not (Figures 3, S5). Using metagenomic classifications sediment depth was the primary driver in community composition (PERMANOVA; pseudo-F=13.91; R^2^=0.39, p=0.001;) with site the secondary driver in community composition (PERMANOVA; pseudo-F=11.86; R^2^=0.08, p=0.001). Community composition was not significantly affected by methane exposure (PERMANOVA; pseudo-F=0.6; p=0.583), or oxygen exposure (PERMANOVA; pseudo-F=0.28; p=0.852).

**Figure 3.**
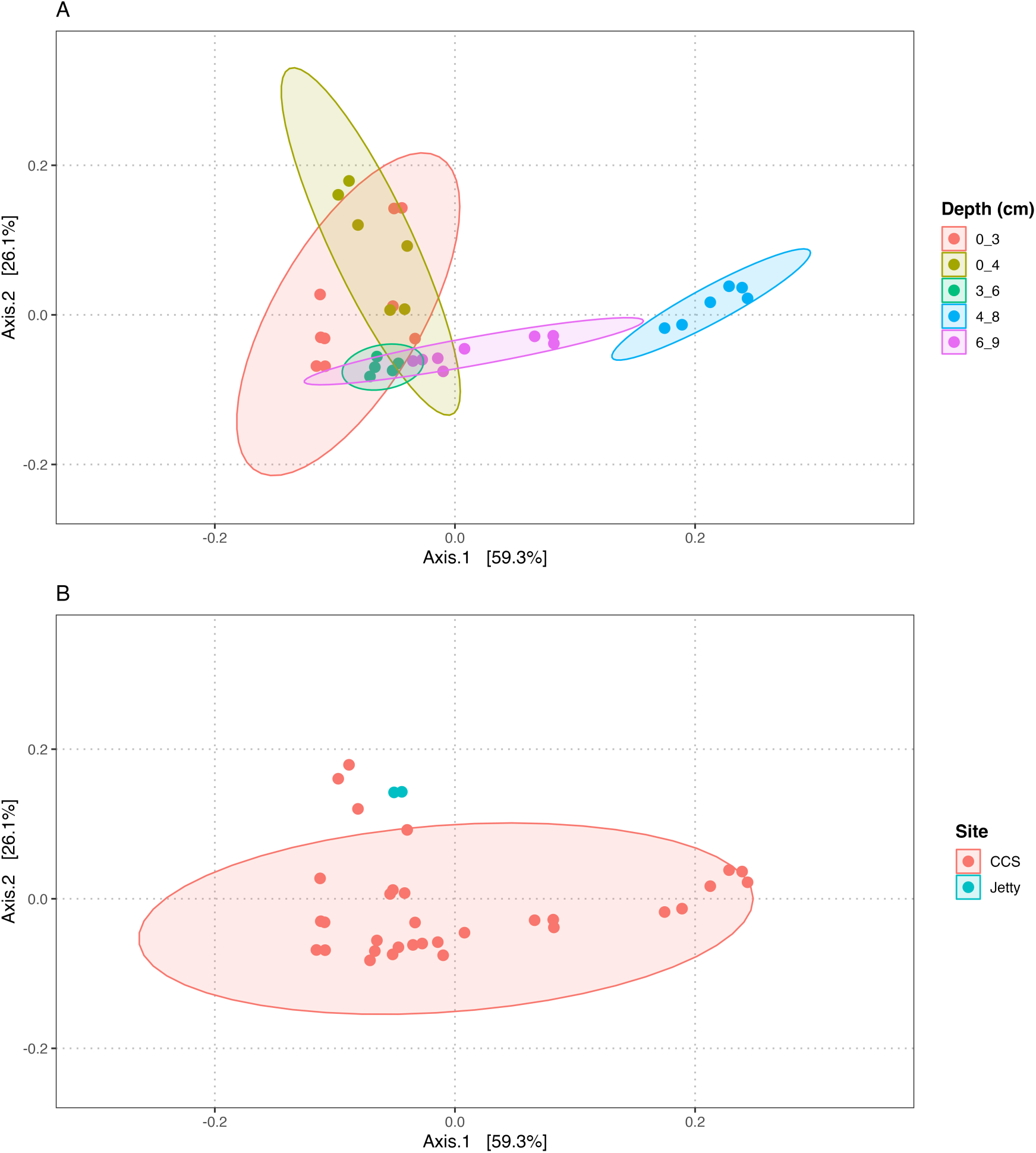
Sediment depth was the most deterministic driver of community composition. Differently colored ellipses represent 95% confidence intervals, and are included as visual aids but do not represent statistical significance. Plots include the effects of Depth (A), and site (B) on community composition.

### Comparison of top bacterial and archaeal taxa

A total of 518M reads were classified, of which 99% were assigned to Bacteria (3,682 genera) and 1% assigned to Archaea (175 genera; Figure 4). The top 15-most abundant genera in each domain accounted for 14.4% and 57.4% of classified reads, respectively. The 15 most abundant bacterial genera were dominated by sulfur cyclers. Sulfate-reducing genera included Desulfogranum (0.91% ± 0.61%), Desulfomarina (0.79% ± 0.42%), Desulfonema (1.12% ± 1.53%), Desulfopila (3.24% ± 1.44%), Desulforhopalus (3.25% ± 1.59%), Desulfosarcina (3.83% ± 3.48%), and Desulfosediminicola (0.97% ± 0.32%). The sulfur-cycling bacteria Sulfitobacter (0.87% ± 0.42%) was present (Sanz-Sáez et al. 2023), along with the sulfur-driven denitrifier Sulfurovum (1.07% ± 0.86%). Wenzhouxiangella (0.95% ± 0.48%), a likely denitrifier (Chunyi et al. 2024) was present as well. Woeseia, also a likely nitrogen cycler, was present (2.45% ± 1.29%), (Mußmann et al. 2017). The top 15 archaeal genera made up ∼30-75 percent of classified archaeal reads (Figure 4B). The majority of top archaea were methanogens, with most reads assigned to the obligate methylotrophic genus Methanococcoides (0.26% ± 0.61%). The non-methanogenic archaeon Archaeoglobus (0.01% ± 0.02%) was present, which reportedly carries out sulfate reduction in certain taxa (Mitchell et al. 2009). Thermococcus, which has been shown as a H_2_ donor for methanogens was present as well. Nitrosopumilus, an ammonia-oxidizing archaeon (Qin et al. 2020) was present in all samples (0.08% ± 0.02%), and was particularly abundant in the upper layer sediments (Figure 4B). Anaerobic methanotrophs were among the top 15 archaea: *Ca. Methanoperedens* (previously ANME-2d), known for coupling nitrate reduction to methane oxidation (Guerrero-Cruz et al. 2021), was present throughout samples (Fig 4B).

**Figure 4.**
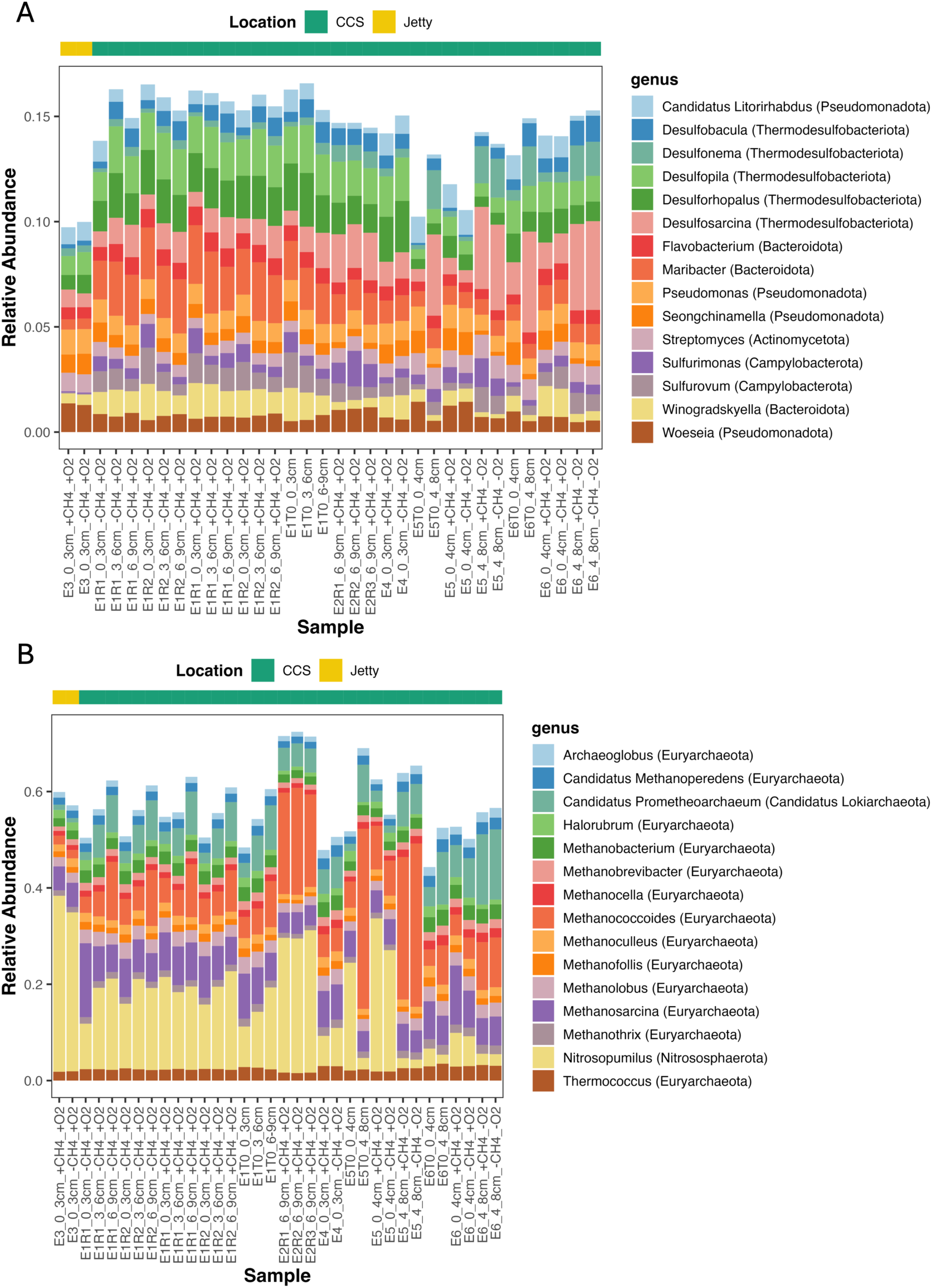
The sediment microbial community was dominated by biogeochemically relevant bacterial and archaeal taxa. The top 15 bacterial genera (A) and top 15 archaeal genera (B) of each sample analyzed by metagenomic analysis. The legend for each plot maps to the genus followed by the corresponding phylum in parentheses.

### Comparison of top methanotrophs

By using metagenomic taxonomy, a community of aerobic and denitrifying anaerobic methane oxidizers was resolved (Figure 5). Across all samples, aerobic methanotrophs were present, including genera belonging to *Gammaproteobacteria*: primarily *Methylobacter* (0.14% ± 0.02%), *Methylomonas* (0.15% ± 0.03%), *Methylomarinum* (0.06% ± 0.01%), and *Methyloprofundus* (0.06% ± 0.01%). Methanotrophs belonging to *Alphaproteobacteria* were present as well, at a lower relative abundance, with the most prominent being *Methylocystis* (0.02% ± 0.004%). The acidophilic *Verrucomicrobiota Methylacidiphilum* was present in low numbers as well (<0.01%). Alongside the nitrate reducing *Candidatus Methanoperedens,* the anaerobe *Candidatus Methylomirabilis* was present across all samples (0.02% ± 0.001%), capable of anaerobically oxidizing methane using nitrite as an electron acceptor (Ettwig et al. 2010).

**Figure 5.**
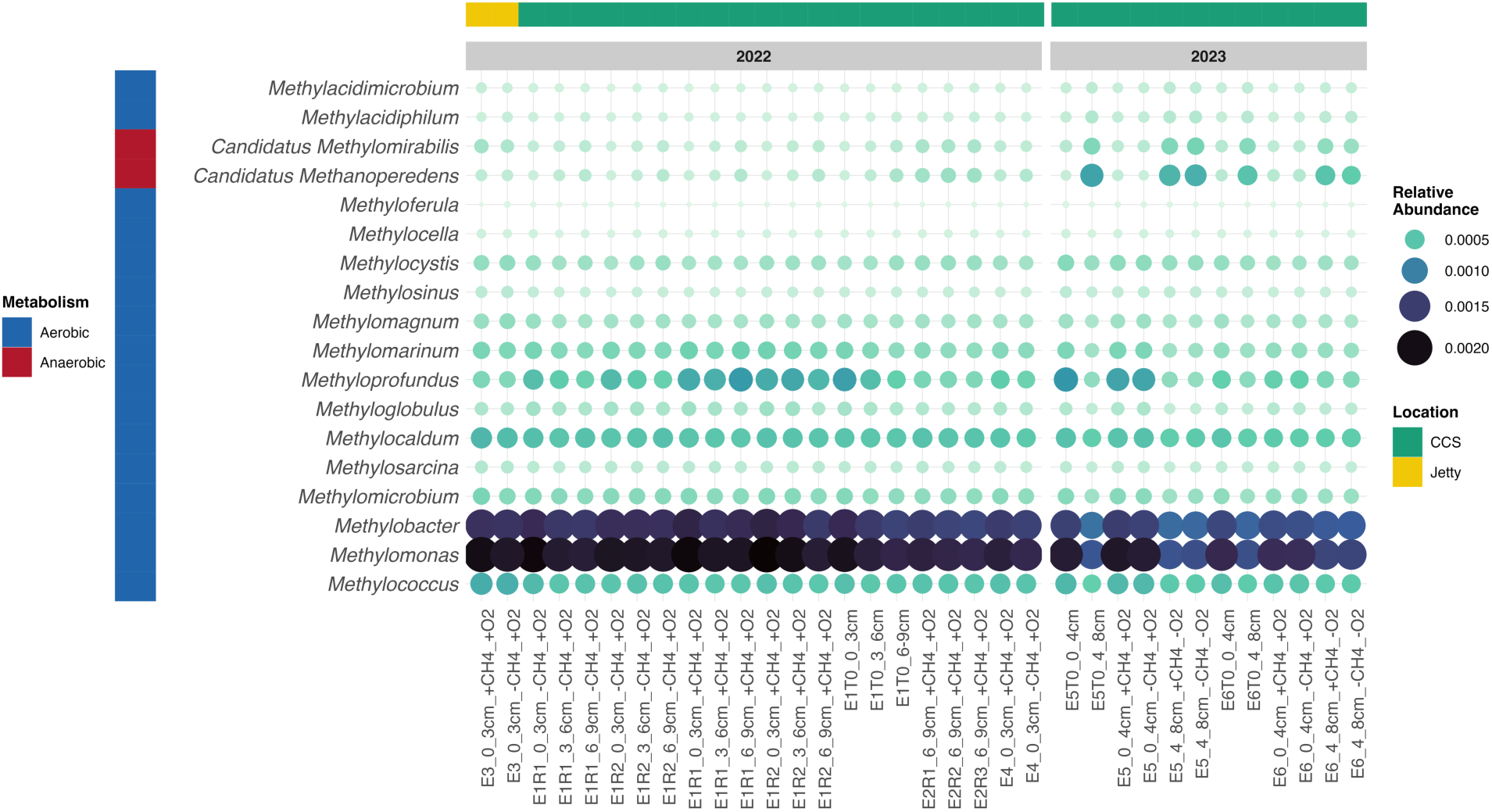
Metagenomic taxonomy reveals a wide diversity of methane oxidizers. Relative abundance of recognized methane oxidizing genera annotated on metagenomic datasets. Relative abundance is represented as lighter color as well as a larger bubble.

### Denitrification-mediated methanotrophs were correlated positively across samples

*Ca. Methylomirabilis* and *Candidatus Methanoperedens* followed congruent abundance patterns, with a significant positive correlation across samples (Figure 6; Spearman’s rho=0.80; p=4.45×10^-7^). The abundance of both taxa increased with depth across both years (Figure 6A). In 2023, 4-8 cm sediment contained 2.18 fold and 4.54 fold higher mean abundance of *Ca. Methylomirabilis* and *Ca. Methanoperedens*, respectively compared to the overlying 0-4 cm sediments. Increased abundance with depth was more modest in 2022, with a 1.74 fold mean increase by *Ca. Methanoperedens* from 0-3 cm to 6-9 cm. *Ca. Methylomirabilis* increased 1.28 fold from 0-3 cm to 6-9 cm, however decreased slightly in mean abundance from 0-3 cm to 3-6 cm (7% decrease). Interestingly, the two Jetty samples yielded the highest residuals from the fitted linear model (Figure 6B; residuals: 8.6×10^-5^ and 5.14×10^-5^) compared to the mean residual value across all points (1.9×10^-5^). The high residual values for the Jetty samples reflect a higher relative abundance of *Ca. Methylomirabilis* in relation to *Ca. Methanoperedens* abundance for this site, but notably both were present.

**Figure 6.**
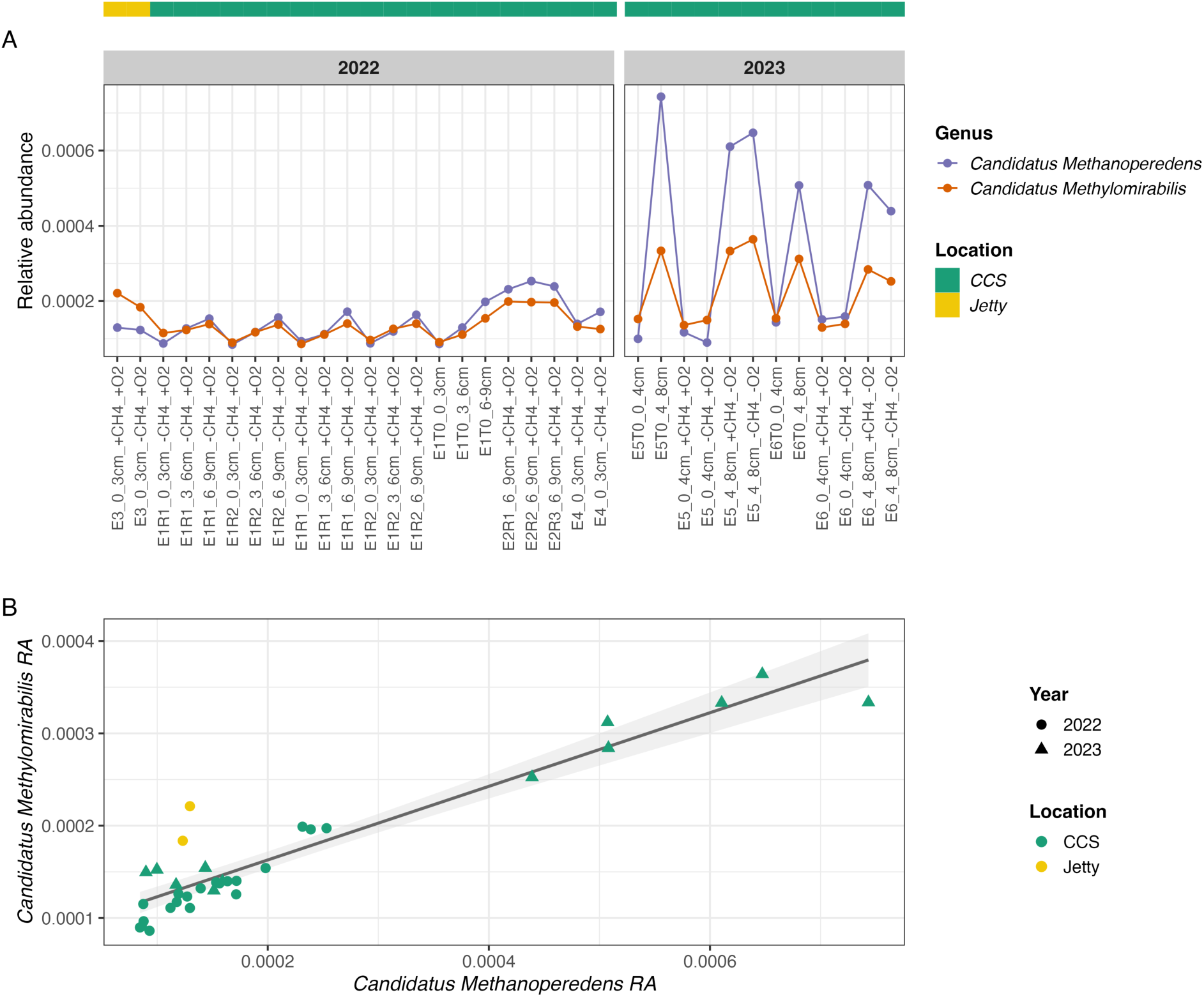
Denitrification-mediated anaerobic methane oxidizers correlated in abundance across samples. The relative abundance of *Ca. Methanoperedens* and *Ca. Methylomirabilis* across samples collected in 2022 and 2023 (A), as well as the correlation between the relative abundance of the two taxa across all samples (B).

### The increased resolution of metagenomic community classification identified more taxa involved in methane cycling

Community composition and taxonomic resolution differed when it was characterized by the V4 region of the 16S rRNA gene compared to metagenomics. 16S rRNA gene analysis identified more bacterial orders (256 metagenomics: 342 16S; Figure 7A) yet metagenomics identified members belonging to more families (667 metagenomics: 501 16S; Figure 7B), and genera (3,539 metagenomics: 711 16S; Figure 7C). Metagenomic approaches classified more archaeal orders (32 metagenomics: 20 16S; Figure 7A) and families (50 metagenomics: 26 16S), with a more than fourfold increase of genera (170 metagenomics: 29 16S). This increase in taxonomic resolution impacted particular groups, including those that are associated with methane oxidation (Figure 7D-F). For example, both metagenomics and 16S identified 5 orders known to contain methane oxidizers (Figure 7D). However, of 11 families involved in methane cycling within our results, 8 were resolved using metagenomics while only 4 families were identified through 16S (Figure 7E). The gap between methods widened at the genus level, with 22 of 33 genera classified by metagenomics, and only one (Methyloprofundus) classified by 16S (Figure 7F). This disparity may be due to common biases such as difficulty amplifying archaea through PCR using standard 16S primers (Raymann et al. 2017; Bahram et al. 2019), database differences (NCBI vs. SILVA), and the distinct classification algorithms underlying each approach (Callahan et al. 2016; Menzel, Ng, and Krogh 2016). We note that with lower levels of classifications, an increasing percentage of unknowns can lead to reduced identified diversity, which can manifest as groups that have methanotrophs as members at the familial level, but none resolved at the genus level. In addition, neither *Ca. Methanoperedens* nor *Ca. Methylomirabilis* were found using 16S rRNA gene analysis at the genus level, and only *Methylomirabilaceae* was found at the family level.

**Figure 7.**
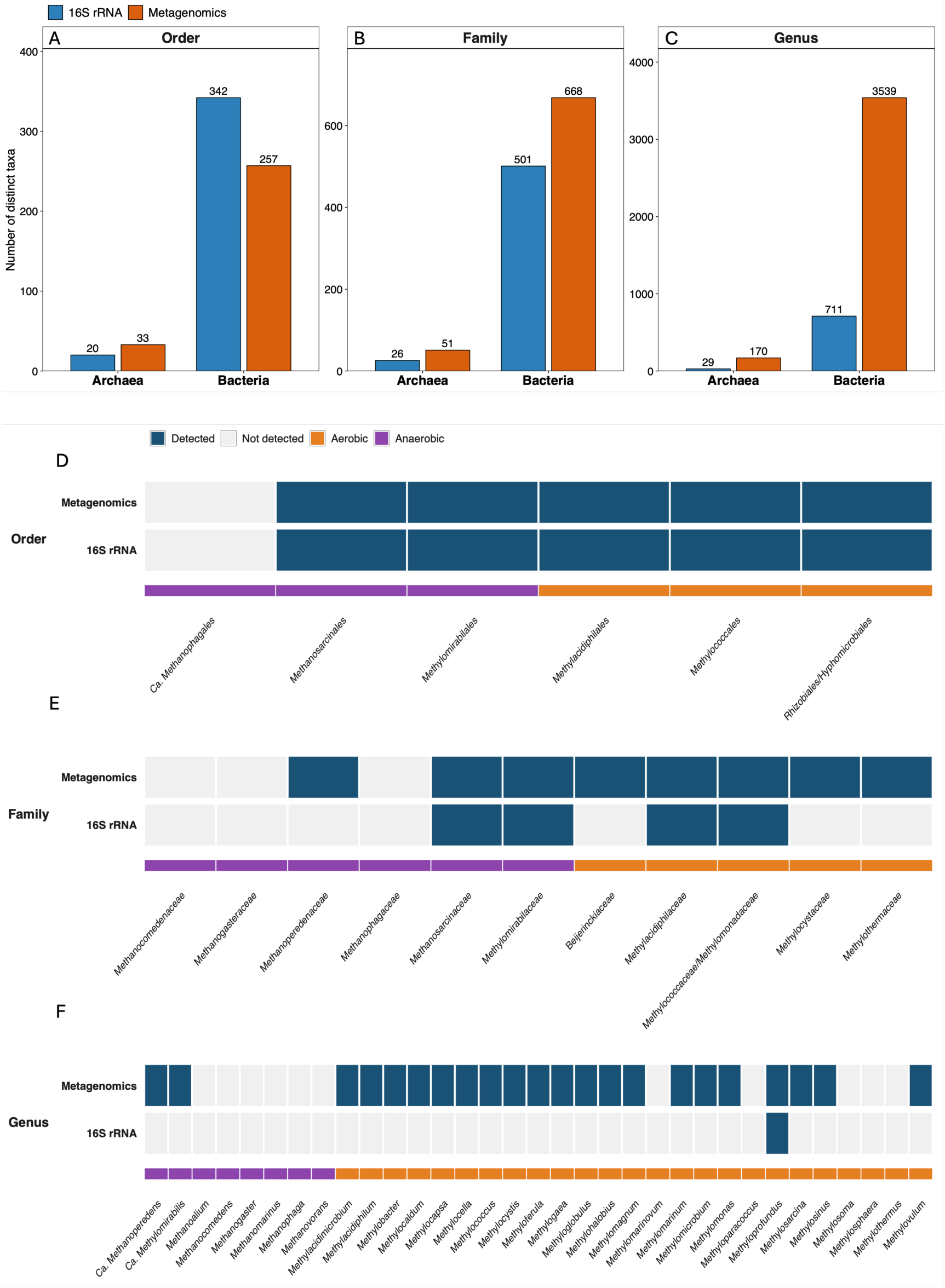
Metagenomics yields higher taxonomic resolution compared to 16S. Comparison of the classification techniques (16S vs. metagenomic short read), both broadly (A-C), and for potential methane oxidizers (D-F). Bar plots represent the number of archaeal or bacterial representatives classified by 16S (blue) or metagenomic techniques (orange) at Order (A), Family (B), and Genus (C) that contain one or more known methane oxidizers. Heatmap plots represent the presence (blue) or absence (grey) of taxa containing one or more methane oxidizers by either metagenomics or 16S at the Order (D), Family (E), or Genus (F) level. Below each heat map, methane oxidizers are identified by their environmental oxygen requirements with either orange (oxic) or purple (anoxic) or orange (oxic) coloring.

### Differential abundance of functional pathways

Genes driving biogeochemical processes related to methane, sulfur, nitrogen, and phosphorus were differentially abundant across samples (Figure 8). Samples clustered strongly by year in which experiments were conducted and the sediment depth (Figure 8). The depth intervals collected in 2023 had particularly strong differences, with 29 out of 38 pathways differentially abundant between 0-4cm and 4-8cm (DESeq2; BH-adjusted p<0.042) with incubations from 4-8 cm clustered functionally with a distinct profile to all other incubations (Figure 8). Incubations from 2023 collected from shallow sediment in oxic incubations, have proportionally more genes coding for denitrification (DESeq2; Log2FC 0.51, p=4.9×10^-10^), and AOM (DESeq2; Log2FC 0.29, p=1.7×10^-4^) in contrast to the deeper 4-8cm samples from those same incubations. Deeper sediments across all experiments (>3cm) have a greater abundance of methanogenesis pathway genes, specifically for the Central Methanogenic Pathway (DESeq2; Log2FC 0.40, p=2.94×10^-6^), and methylotrophic methanogenesis (DESeq2; Log2FC 0.34, p=7.32×10^-5^), reflecting the high diversity of methanogens described above (Figure 4B). Interestingly, samples taken from the Jetty cluster with shallow on-seep samples from E6 taken in 2023 (Figure 8). Oxygen as a treatment was not sufficiently replicated to test its role on the metabolisms present statistically. However, since the dominant changes were between sites and depths, it is unlikely to have impacted our conclusions.

**Figure 8.**
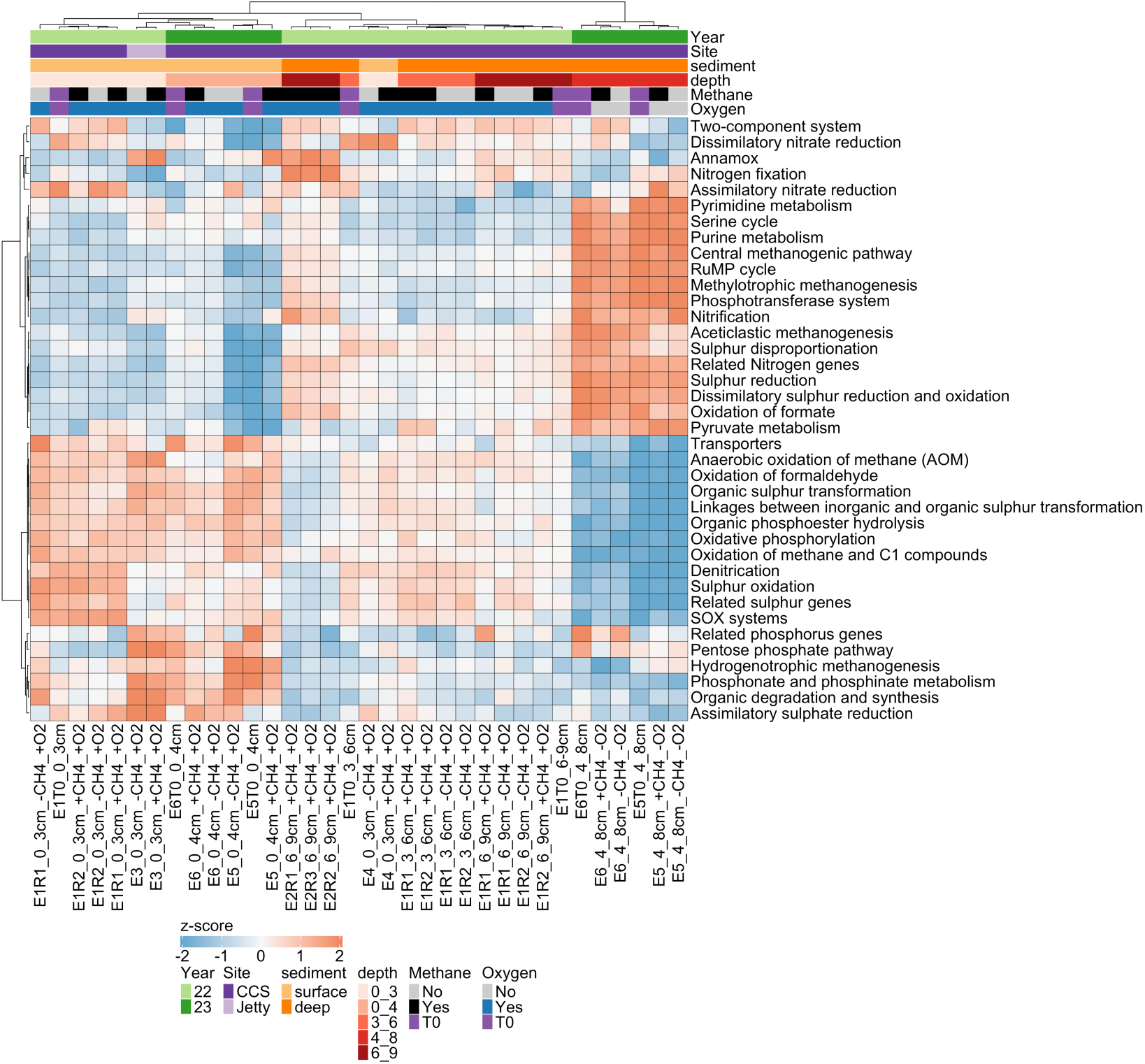
Biogeochemical cycling genes differ in abundance across samples. Prior to visualization, pathway counts were transformed via Variance Stabilizing Transformation using DESeq2, then row-scaled to z-score so that each pathway is centered on its mean abundance across samples. Red boxes indicate pathways that are relatively higher in abundance while blue boxes indicate pathways that are relatively lower in abundance. The distance between samples was calculated using Euclidean distance and using a Ward clustering algorithm.

## Discussion

In Ross Sea sediment, the presence of methane oxidation alongside known methane oxidizers was observed across replicated experiments, years, experimental set-up, and study site, and continued through different oxygen and methane conditions. We expected a clear difference in on-seep vs. off-seep communities and oxidation values when comparing the Jetty samples against the CCS samples based on previous research (Thurber, Seabrook, and Welsh 2020). We found that, although the community shifted across samples, methane oxidation occurred throughout. The Jetty hosted the same methanotrophs present in CCS and oxidized methane, albeit with lower relative abundance of methanotrophs (Figures 4,5). The presence of methane oxidizers suggests that although the microbial community may shift in an emergent seep (Thurber et al. 2020), the taxa capable of methane oxidation may be ubiquitously present in the region, possibly due to the high abundance of methanogens in the sediments even in control areas (Figure 4). While limited to 2 sites within this study, it appears that active methane cycling is a commonality in Antarctic shallow habitats.

While globally, sulfate reduction couples with methane oxidation, this did not appear to be the case at the CCS. Methane oxidation coupled to the electron acceptor sulfate is understood as the primary methane filter in marine sediments and is particularly important in deep-sea methane seeps, reportedly responsible for removing 70-90% of methane in seep sediments before reaching the water column (Caldwell et al. 2008; Knittel and Boetius 2009). However, the only anaerobic methane oxidizers found in our study were denitrification dependent (Figures 4,5,6,7): *Ca. Methanoperedens* and *Ca. Methylomirabilis*. We note that the relative abundance values of both taxa are highly correlated across samples (Figure 6), suggesting tight metabolic cooperation. This has been observed across other systems (Vaksmaa et al. 2017) including through the observation of mixed aggregates of the two taxa under enrichment (Guerrero-Cruz et al. 2018). Given that *Ca. Methanoperedens* produces nitrite as a byproduct, it can likely be co-opted by *Ca. Methylomirabilis* under the right conditions (Guerrero-Cruz et al. 2021). Together, these findings implicate nitrogen cycling as a component of anaerobic methane oxidation at CCS. These findings alter the understanding of the rate at which a biological methane filter takes hold on an Antarctic seep. For example, an early presence of *Ca. Methylomirabilis* and *Ca. Methanoperedens* could mark a faster biological filter than previously predicted, with a potential doubling time of 1-3 weeks and 2-3 weeks, respectively, compared to ANME-1’s reported 7 months (Guerrero-Cruz et al. 2021; L. He et al. 2023; Knittel and Boetius 2009).

The aerobic oxidation of methane was a ubiquitous methane-fueled metabolism. Excess methane that escapes cold-seep sediment is most often oxidized through methane oxidizing bacteria (MOB) in the surface sediment and water column (Leonte et al. 2017; Joung et al. 2022). Increasing evidence suggests that the range of MOBs extends even into anoxic sediments (He et al. 2022; Oswald et al. 2016). For example, He et al. (2022) used stable isotope probing to establish the anoxic assimilation of methane by Gammaproteobacterial methanotrophs in an Arctic lake even as deep as 70cm within sediments. Oswald et al. (2016) established aerobic methanotrophs oxidizing methane in anoxic settings stimulated through the addition of iron and manganese oxides. These findings suggest that MOBs are capable of methane oxidation in undetectable oxygen conditions. In addition, sediment disturbances such as advection (Ruff et al. 2016) and macrofaunal burrowing (Deng et al. 2020; Thurber et al. 2013) can introduce oxygen into sediments. Given that the genera *Methylobacter* and *Methylomonas* are the most abundant methanotrophs across depths in our results (Figure 5), it is likely that they contribute to methane oxidation both within and above the sediments and were in some part responsible for observed methane oxidation in this study (Figure 2). Notably, both taxa were also only resolved with metagenomic community characterization (Figure 7).

Our findings, together with prior studies, contribute to an evolving understanding of nitrogen cycling in methane seeps (Wang et al. 2025; Quan et al. 2024; Jing et al. 2020; Dekas, Poretsky, and Orphan 2009; Semler et al. 2025). Marine cold seeps have been increasingly established as hotspots for nitrogen cycling, including nitrogen loss (Jiang et al. 2025), fixation (Dekas, Poretsky, and Orphan 2009), and transformation (Li et al. 2024). While many of these studies demonstrate community-wide effects on nitrogen cycling, fewer find direct or indirect linkages between methane and nitrogen (Jing et al. 2020; Dekas, Poretsky, and Orphan 2009; Dekas et al. 2014), and those that do primarily discuss the influence on ANME-SRB consortia. This includes nitrogen fixation by ANME-2 followed by sharing with SRBs (Dekas et al. 2014; Dekas, Poretsky, and Orphan 2009; Metcalfe et al. 2021), as well as nitrate availability structuring SRB partner choice (Green-Saxena et al. 2014). In contrast, we found evidence of denitrification as a direct link to marine methane oxidation (Figures 5,6,7). The presence of denitrification-mediated anaerobic methane oxidation (dAMO) has been well established in terrestrial and freshwater ecosystems (Wei et al. 2022; Guerrero-Cruz et al. 2021; M. Zhang et al. 2024), and we expand this finding to marine Antarctic Seeps. These findings bolster similar discoveries that dAMO is a methane sink in marine coastal (Shen et al. 2016), and cold-seep environments (Jing et al. 2020). Antarctic cold seeps occupy an unusual position in which they undergo seasonal productivity and nutrient flux as a coastal ecosystem, yet exist under consistent low temperature and prolonged periods without light. Given the unique characteristics of this system, it is likely that further study in the Ross Sea will further contribute to the understanding of dAMO in coastal and cold seep environments across the globe.

Given the vast amounts of predicted methane in the Antarctic (Wadham et al. 2012) and emergent seepage (Seabrook et al. 2025), understanding the microbial community in the region informs current and future carbon regimes. Microbial succession of the seep continues to shift from reported ANME-1 in 2016 (Thurber et al. 2020) to dAMO driven taxa alongside various aerobic methanotrophs in 2022 and 2023 (Figure 7). Given the putative link between nitrogen and methane oxidation, alongside seasonal shifts in regional nitrogen concentration (Barry 1988), we posit that methane oxidation rates are likely dynamic in the region and depend on season and nutrient availability. Our findings supplement the evidence that methane seep systems are heterogeneous at both local and global scales (Seabrook et al. 2018; Ruff et al. 2015; Semler and Dekas 2025), and underscore the importance of studying the similarities and differences of Antarctic methane seeps. Continued examination of these systems aids in our understanding of what appears to be an emergent phenomenon with potentially global impacts on a crucial greenhouse gas.

## Conclusions

With the recent emergence of methane seeps across the Ross Sea, it is crucial to understand the capabilities of their microbiomes as a methane filter and their impact on biogeochemical cycling in the region. Given the vast amounts of predicted methane reserves in the Antarctic ice sheet and the discovery of dozens of emergent seeps within the Ross Sea, our findings increase the understanding of microbial capability and potential as it relates to methane oxidation in the region. Given that an ANME-SRB consortium was not present in either 16S or metagenomic analysis, methane cycling appeared to be driven by denitrification-mediated anaerobic oxidation of methane and aerobic methanotrophy. Both methane oxidation and community composition also highlight the ubiquity of methane cycling in the shallow habitats of the Ross Sea.

## Acknowledgements

We would like to extend our thanks to the Center for Quantitative Life Sciences (CQLS) at Oregon State University for providing the sequencing infrastructure and computing resources necessary to complete this project. Use of Artificial intelligence software We would also like to thank Lila Ardor Bellucci for contributing to the field work necessary for this project. We remain indebted to the support of the USAP Divers Rob Robbins, Steve Rupp, and Alex Brett as well as the McMurdo community who made this research possible.

## Notes

https://doi.org/10.5281/zenodo.21157354

https://github.com/jacob8776/antarctic_seep_incubations_peerJ

### Competing Interest Statement

The authors have declared no competing interest.

### Funding

- U.S. National Science Foundation: 2425835, 2535603, 2325046, 2139319

