## Supplemental figures and extended legends for "When seeps give ANME-SRB the cold shoulder: putative role of denitrification mediated methane oxidation in an Antarctic Cold Seep"

### Supplemental material

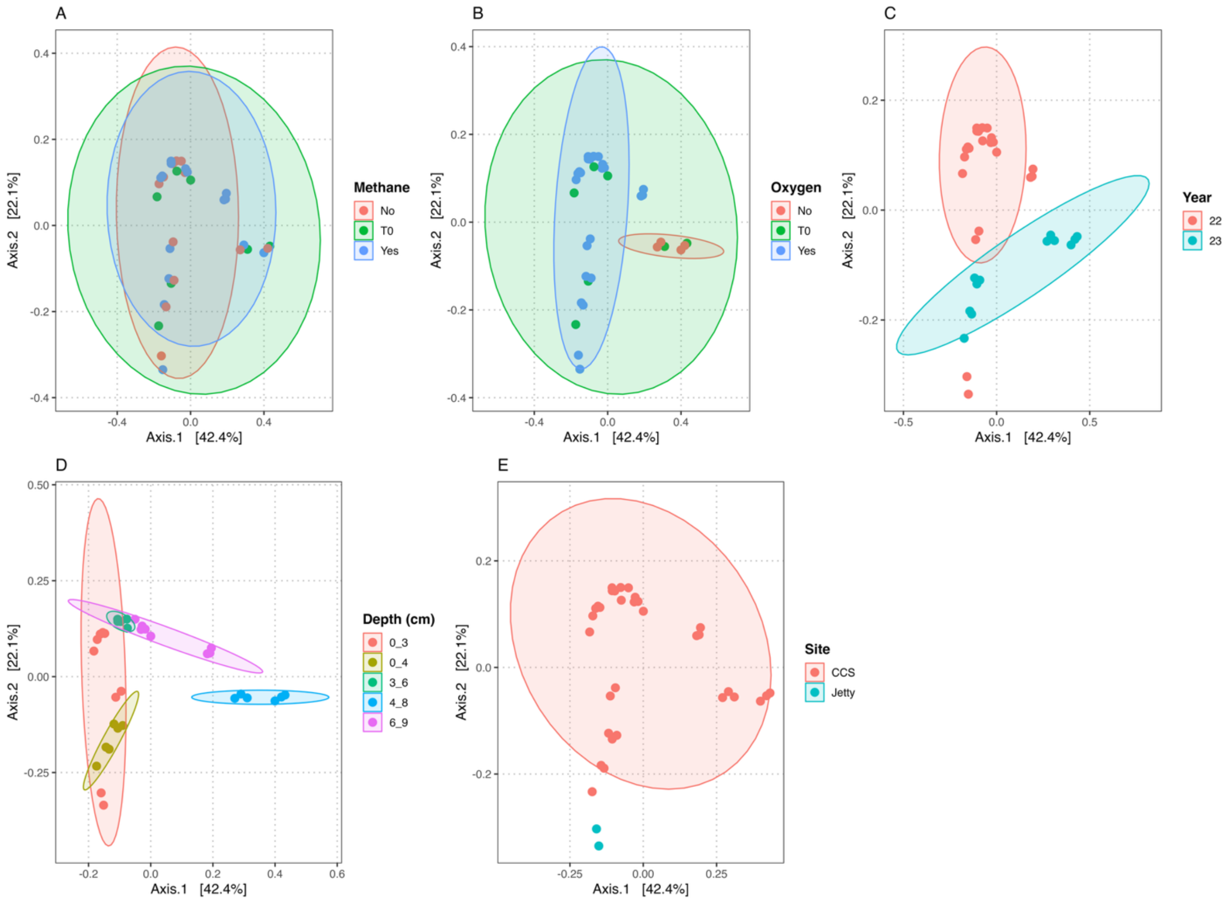


**Figure S1. PCoA of 16S taxonomy conglomerated at the genus level across samples.** Differently colored ellipses represent 95% confidence intervals, and are included as visual aids but do not represent statistical significance. Plots include the effects of Methane (A), Oxygen (B), Year (c), depth (D), and site (E) on community composition.

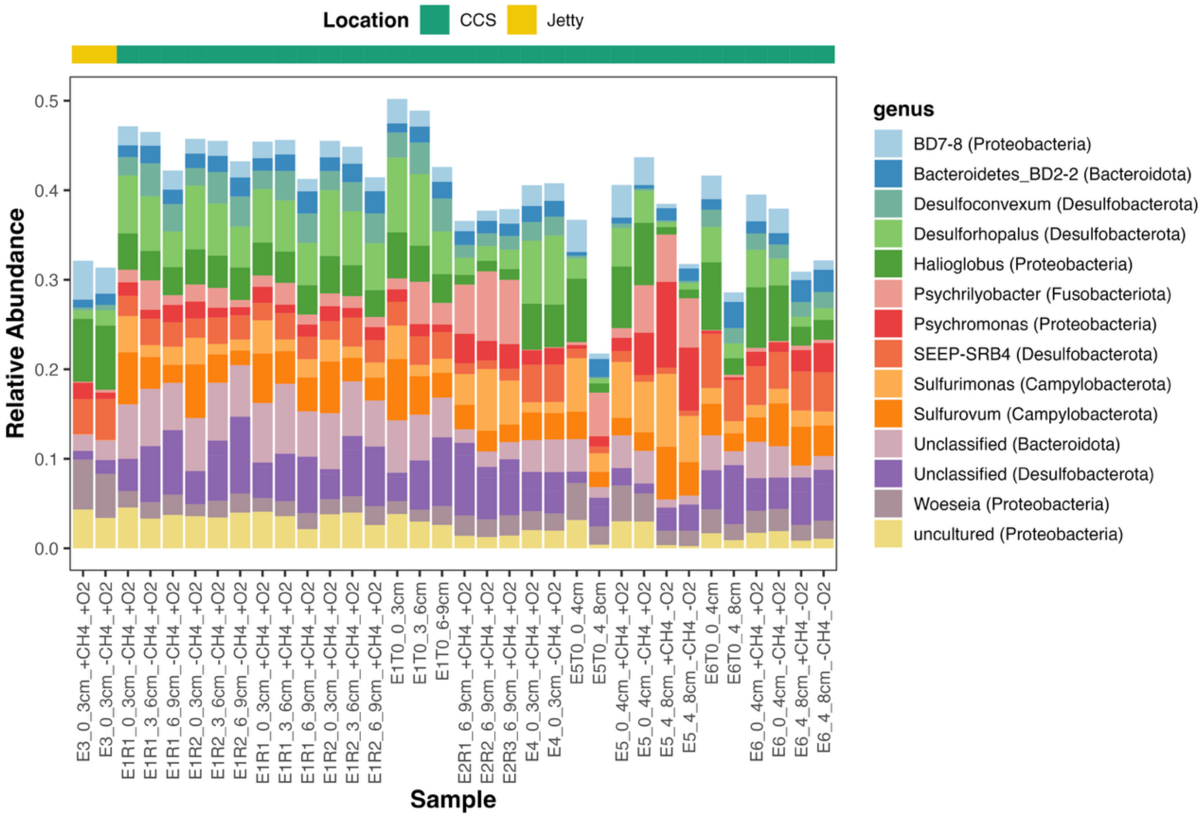


**Figure S2. The top 15 bacterial genera of each sample analyzed by 16S rRNA gene analysis.** The legend for each plot maps to the genus followed by the corresponding phylum in parentheses for each genera. Annotations were made according to the taxonomic schemes in SILVA 138.1 database. Data was normalized prior to the construction of this plot by using relative abundance, dividing reads from each taxon within a sample with the total number of bacterial reads for the sample.

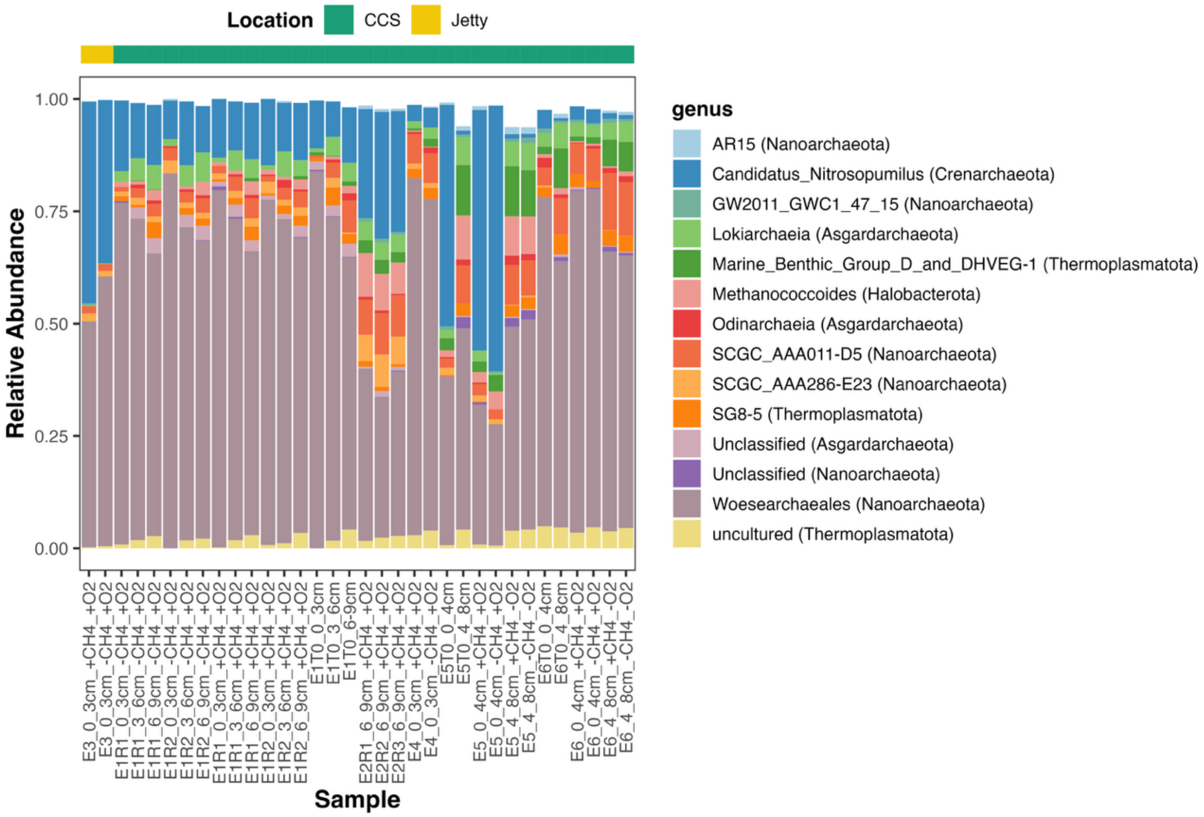


**Figure S3. The top 15 archaeal genera of each sample analyzed by 16S rRNA gene analysis.** The legend for each plot maps to the genus followed by the corresponding phylum in parentheses for each genera. Annotations were made according to the taxonomic schemes in SILVA 138.1 database. Data was normalized prior to the construction of this plot by using relative abundance, dividing reads from each taxon within a sample with the total number of archaeal reads for the sample.

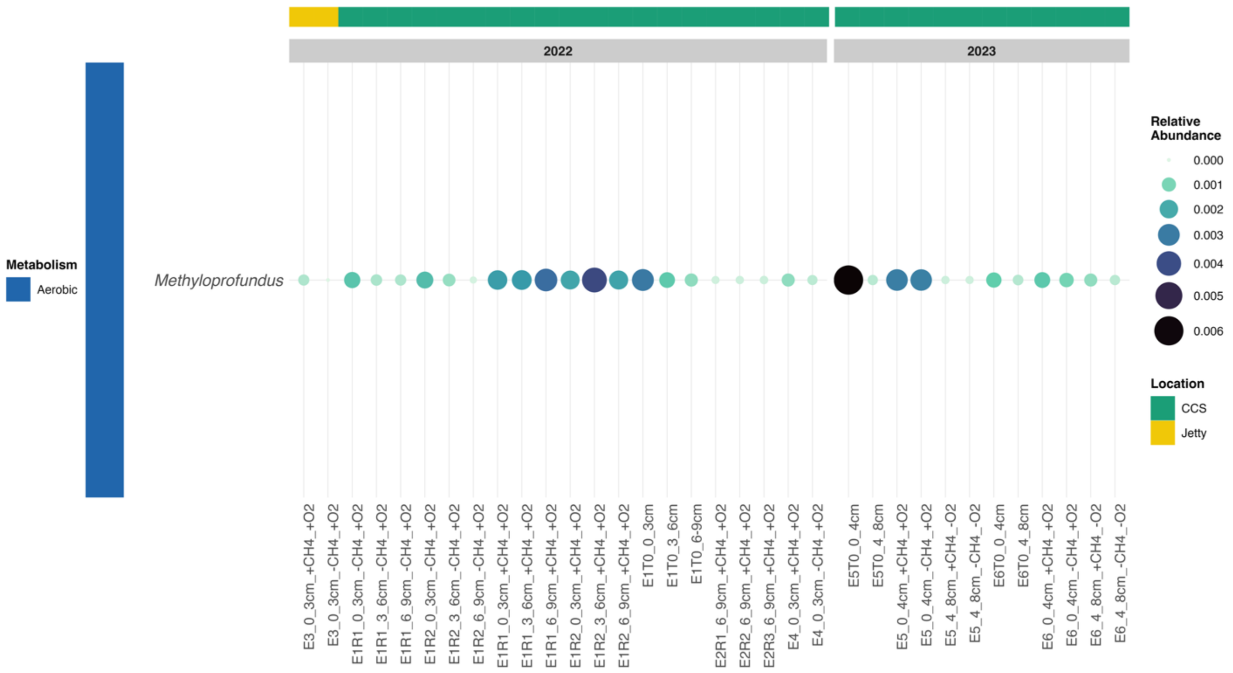


**Figure S4. Relative abundance of recognized methane oxidizing genera annotated on 16S datasets.** Relative abundance is represented as a lighter color as well as a larger bubble.

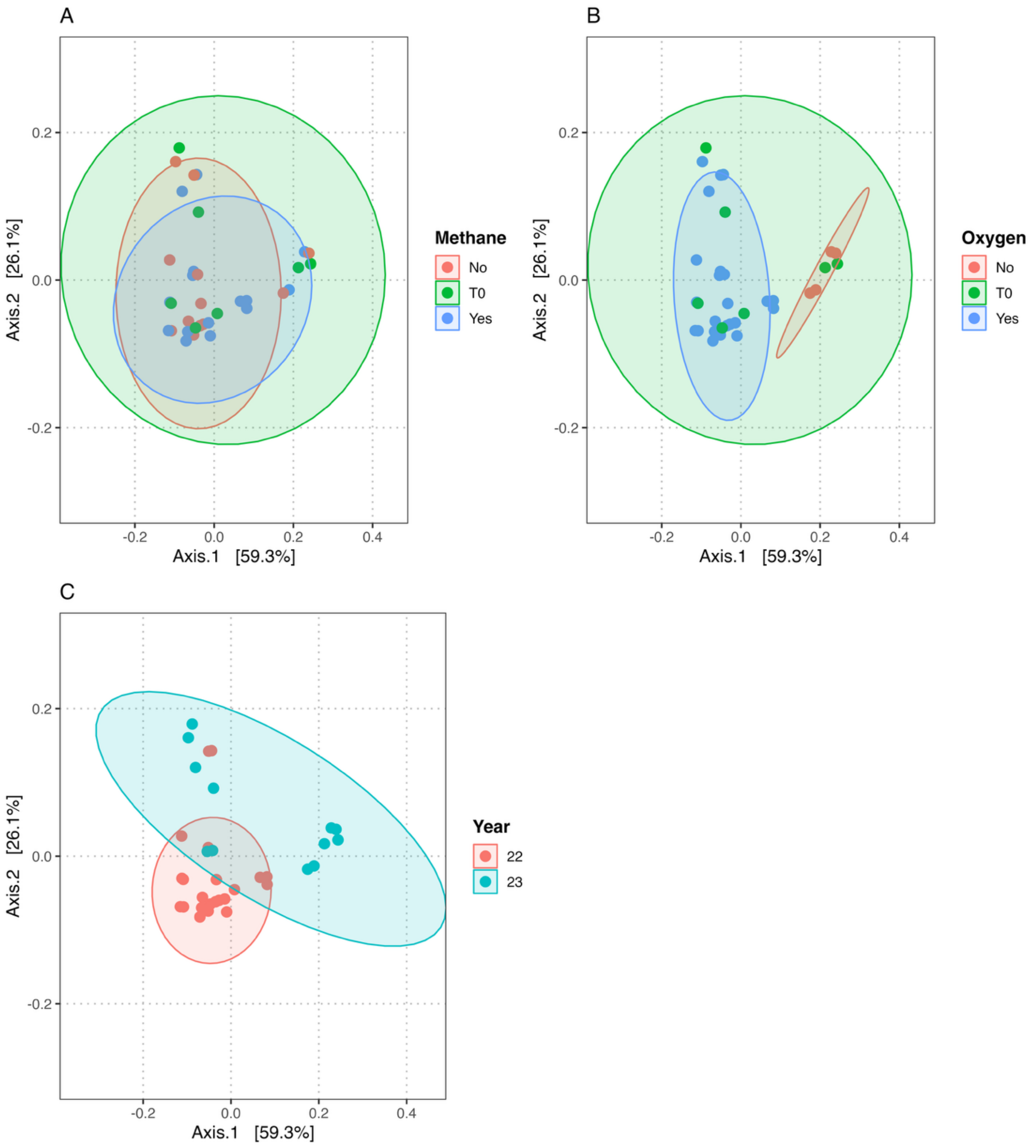


**Figure S5. PCoA of metagenomic taxonomy conglomerated at the genus level across samples.** Differently colored ellipses represent 95% confidence intervals, and are included as visual aids but do not represent statistical significance. Plots include the effects of Methane (A), Oxygen (B), and Year (C) on community composition.

**Table S1. 2022 and 2023 Experiment methods.** Columns including the sample collection site, sediment depths at which slurries were made, days incubated, and whether an abiotic control was included in the experiment. It also specifies whether any part of the incubation showed success in maintaining anoxia.

**Table S2. Oxygen values recorded within incubation vials at the breakdown of each experiment.**

**Table S3. Change in $^{13}C_{dic}$ across experiments in 2022 and 2023.** These values also include the filtered sea water controls (FSWC) with no methane oxidation observed.

**Table S4. Taxonomic reconciliation between SILVA and NCBI databases for methanotroph taxa.**

**Table S5. Taxonomy set for each methanotrophic genus in the study, reconciled between databases.**

**Table S6. Statistics of reads assigned to methanotroph genera across 16S and metagenomics methods.** This table includes the total reads assigned to each genus across all samples, the mean reads across samples, max reads in one sample, and the number of samples that a given genus was detected.

**Table S7. Number of reads and relative abundance of methanotroph genera across 16S and metagenomic methods, by sample.**

**Dataset S1. Forward and reverse primers used for 16S amplification.** PCR primers used to generate the 16S raw data for the study.
